# *Snca*-GFP knock-in mice reflect patterns of endogenous expression and pathological seeding

**DOI:** 10.1101/2020.05.05.077321

**Authors:** Anna Caputo, Yuling Liang, Tobias D. Raabe, Angela Lo, Mian Horvath, Bin Zhang, Hannah J. Brown, Anna Stieber, Kelvin C. Luk

## Abstract

Alpha-synuclein (aSyn) participates in synaptic vesicle trafficking and synaptic transmission, but its misfolding is also strongly implicated in Parkinson’s disease (PD) and other neurodegenerative disorders known as synucleinopathies where misfolded aSyn accumulates in different regions of the central and peripheral nervous systems. Although increased aSyn expression levels or altered aggregation propensities likely underlie familial PD with SNCA amplification or mutations, the majority of synucleinopathies arise sporadically, indicating that disease can develop under normal levels of wildtype aSyn. We report here the development and characterization of a mouse line expressing an aSyn-GFP fusion protein under the control of native *Snca* regulatory elements. Regional and subcellular localization of the aSyn-GFP fusion protein in brains and peripheral tissues of knock-in (KI) mice are indistinguishable from that of wildtype littermates. Importantly, similar to wildtype aSyn, aSyn-GFP disperses from synaptic vesicles upon membrane depolarization, indicating that the tag does not alter normal aSyn dynamics at synapses. In addition, intracerebral injection of aSyn pre-formed fibrils into KI mice induced the formation of aSyn-GFP inclusions with a distribution pattern similar to that observed in wildtype mice, albeit with attenuated kinetics due to the GFP tag. We anticipate that this new mouse model will facilitate *in vitro* and *in vivo* studies requiring *in situ* detection of endogenous aSyn, therefore providing new insights into aSyn function in health and disease.

**Significance Statement:** Alpha-synuclein (aSyn) participates in synaptic vesicle function and represents a major component of the Lewy pathology found in Parkinson’s and related neurodegenerative diseases. The function of aSyn and the sequence of events leading to its aggregation and neurotoxicity are not fully understood. Here we present a new mouse model in which Enhanced Green Fluorescence Protein (GFP) has been knocked-in at the C-terminal of the *Snca* gene. The resulting fusion protein shows identical expression and localization to that of wildtype animals, is functional, and is incorporated into pathological aggregates *in vitro* and *in vivo*. This new tool allows for monitoring aSyn under a variety of physiological and pathological conditions, and may uncover additional insights into its function and dysfunction.

## Introduction

Alpha-synuclein (aSyn) is a protein prominently expressed in neurons and enriched at presynapses (Maroteaux et al., 1988). Although its precise functions are not fully understood, collective evidence suggests that aSyn regulates synaptic vesicle trafficking and fusion (Burre et al., 2018; Logan et al., 2017; Nemani et al., 2010; Wang et al., 2014). aSyn also constitutes the major component of Lewy bodies and Lewy neurites, intraneuronal inclusions characteristic of Parkinson’s and a group of neurodegenerative diseases known as synucleinopathies (Bendor et al., 2013; Burre et al., 2018; Goedert et al., 2013). Although mutations or amplification of the aSyn gene were linked to familial PD (Koros et al., 2017; Kruger et al., 1998; Polymeropoulos et al., 1997; Schneider and Alcalay, 2017; Singleton et al., 2003; Zarranz et al., 2004) the majority of PD cases are sporadic and develop in the presence of normal levels of wildtype (wt) aSyn.

Several aSyn animal models have been generated to study aSyn function and pathobiology(Visanji et al., 2016). Accumulation of aggregated aSyn, loss of dopaminergic neurons, and behavioral impairments have been reported in multiple lines. However, the majority of these models rely on the overexpression of human aSyn, often bearing disease-associated mutations, under the control of a heterologous (i.e. non-Snca) promoter, and in the presence of endogenous mouse aSyn expression (Emmer et al., 2011; Giasson et al., 2002; Kahle et al., 2001; Masliah et al., 2000; Matsuoka et al., 2001; Tofaris et al., 2006; van der Putten et al., 2000). These features might confound data interpretation, especially in relation to sporadic PD. These models also poorly recapitulate the prion-like propagation of pathological aSyn suggested by the staged distribution of aSyn deposits in diseased human brains (Braak et al., 2003; Goedert et al., 2013; Jucker and Walker, 2018). Recent studies demonstrating that aSyn pathology formation and spread can be initiated in wt mice following inoculation of recombinant aSyn preformed fibrils (PFFs) (Luk et al., 2012; Masuda-Suzukake et al., 2014; Rey et al., 2018) or human brain derived aSyn aggregates (Masuda-Suzukake et al., 2013; Peng et al., 2018; Recasens et al., 2014) further support this hypothesis, thus enabling the modeling of these processes without aSyn ectopic expression. Exposure to PFFs also induces aSyn pathology in primary neurons, allowing for cellular and molecular characterizations (Volpicelli-Daley et al., 2011). These models provide additional structural and temporal resolution for observing aSyn pathogenesis, yet they do not allow to monitor aSyn in real-time. To this aim, two mouse models overexpressing human aSyn tagged with GFP have been reported (Hansen et al., 2013; Rockenstein et al., 2005). Transient overexpression of aSyn-GFP has also been used in primary neurons (Fortin et al., 2005; McLean et al., 2001; Volpicelli-Daley et al., 2014). These models have provided valuable insights about the trafficking of aSyn at synapses (Fortin et al., 2005), aSyn aggregate formation and development (Osterberg et al., 2015; Unni et al., 2010) and their link to synaptic/axonal dysfunction (Scott et al., 2010; Volpicelli-Daley et al., 2014) and cell death (Osterberg et al., 2015).

However, aSyn is differentially expressed in specific neuron subtypes (Hawrylycz et al., 2012; Taguchi et al., 2016; Uhlen et al., 2015) and its function and propensity to aggregate is dependent on expression levels (Logan et al., 2017; Luna et al., 2018; Nemani et al., 2010; Scott and Roy, 2012), thus making the use of the endogenous promoter highly desirable. We therefore created a mouse in which GFP was knocked-in at the carboxy-terminal of the endogenous Snca gene. The resulting mouse displays aSyn-GFP expression at wt levels, only in neurons and with the same distribution as wt aSyn. Moreover, the fusion-protein localizes correctly to synaptic vesicle and participates in the synaptic vesicle cycle. Importantly, aSyn-GFP is incorporated in Lewy-like pathology seeded by exposure to PFFs. We anticipate that this new tool will allow for further studies aimed at better understanding aSyn physiology and pathobiology.

## Results

### *Snca-GFP* knock-in mice express wildtype levels of aSyn-GFP fusion protein

We generated a novel mouse line that genetically encodes fluorescent aSyn by using homologous recombination to insert the cDNA for enhanced-GFP into the *Snca* locus of murine embryonic stem cells (Figure 1A; see also *Methods*). The construct was targeted to the 3’ end of exon 6 while keeping the surrounding endogenous regulatory elements intact. The modified *Snca-GFP* gene was predicted to transcribe a fusion protein consisting of wild type (wt) murine aSyn with a C-terminal GFP tag. Embryonic stem cells containing the recombined construct following electroporation were confirmed by Southern blot analysis (Figure 1A, right panel) and a validated clone (1B6) was implanted in C67/B6 blastocysts to yield founder lines. Two founder mouse lines were derived and offspring with normal karyotypes were further expanded by breeding out onto a C57/Bl6J background. PCR genotyping with primers targeting exon-intron boundaries revealed mice with either wt, heterozygous (*Snca*^*wt/GFP*^), or homozygous (*Snca*^*GFP/GFP*^) knock-in genotypes (Figure 1B). Both knock-in genotypes were fertile and did not show detectable differences in lifespan nor any overt behavioral defects up to two years of age.

**Figure 1.**
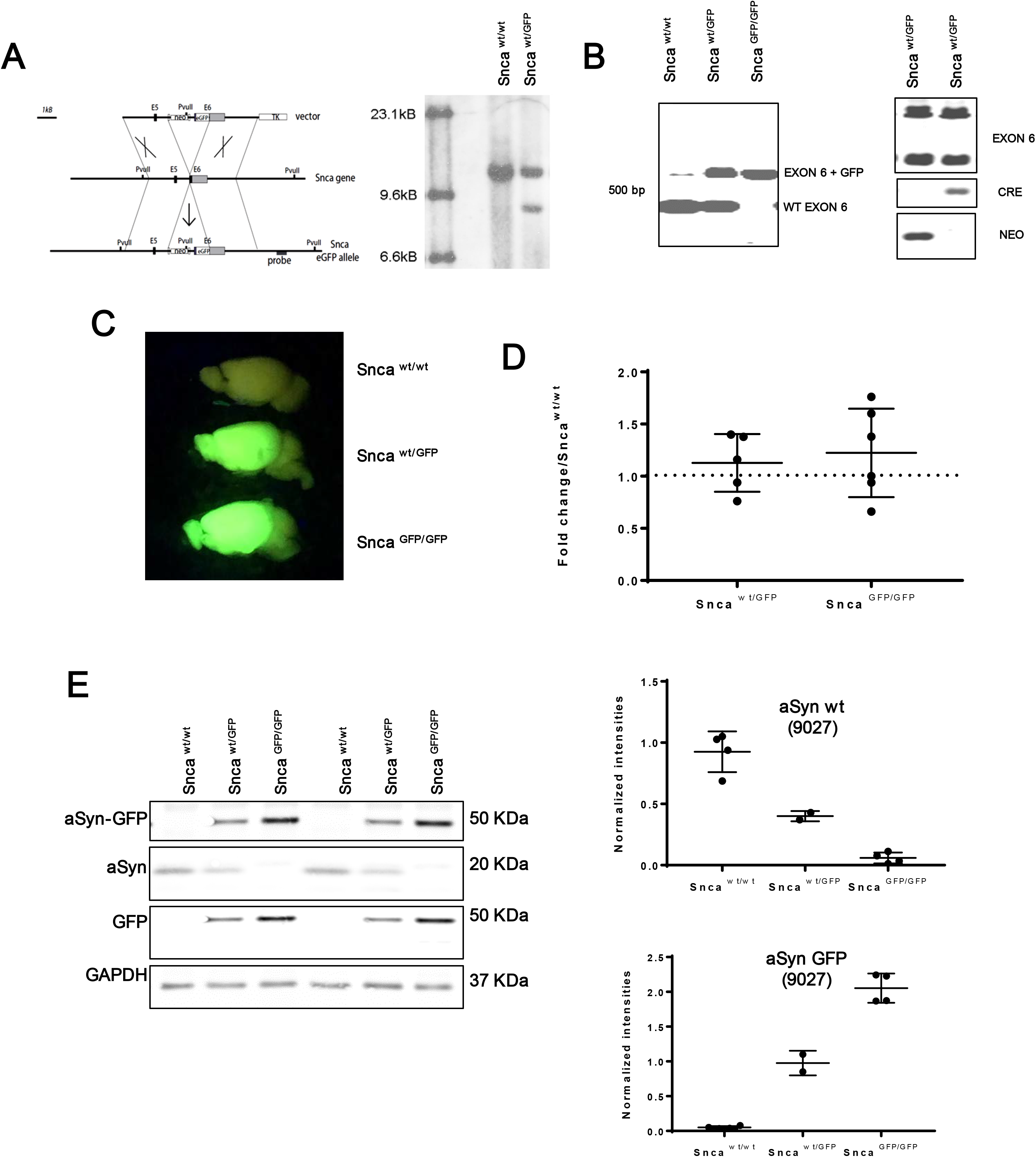
Design and validation of *Snca*-GFP KI mouse. **A)** The GFP sequence was knocked in at the carboxy-terminal end (Exon 6; E6) of the Snca gene, resulting in the expression of aSyn-GFP under the endogenous regulatory elements (see Methods for additional details). Right panel shows Southern blot of geneticin-resistant ES cells electroporated with the targeting vector in A following enzymatic digestion with the restriction enzyme PvuII. Note the additional PvuII restriction site introduced by the integration of the targeting vector. **B)** PCR amplification of wt and Snca^GFP^ from hetero- and homozygous mice. Exon 6 comprises the fusion region. **C)** Brains from heterozygous and homozygous Snca^GFP^ mice fluoresce when illuminated under blue light. **D)** Quantification of aSyn mRNA levels in brain homogenates of hetero- or homozygous KI mice compared to wt littermates. Data are expressed as fold-change in aSyn normalized to actin. Mean and SD values are shown (N= 5-6 per condition). No statistically significant difference was found between groups using One-Way ANOVA (Kruskal-Wallis test and Dunn’s multiple comparison test). **E)** Levels of aSyn and aSyn-GFP in brain homogenate from wt, heterozygous and homozygous mice. Graphs show quantification of the immunoblot data expressed as intensities of aSyn (Syn9027) or aSyn-GFP normalized to GAPDH levels. Mean and SEM values shown (N= 2-4 per group).

Upon illumination with blue light, brains and spinal cords from *Snca*^*wt/GFP*^ and *Snca*^*GFP/GFP*^ mice fluoresced brightly, indicating expression of the aSyn-GFP fusion protein in tissues where aSyn is normally abundant (Figure 1C). Quantification of aSyn mRNA by RT-PCR showed that total transcript levels did not differ between KI and wt mice (Figure 1D). We further confirmed that this represented the expression of an intact aSyn-GFP fusion protein by subjecting brain homogenates derived from wt and KI mice to immunoblot analysis. Probing with antibodies against aSyn and GFP showed a 45-kDa product corresponding to the predicted molecular weight of aSyn-GFP in *Snca* ^*GFP/GFP*^ and *Snca*^*wt/GFP*^ littermates that was absent from wt controls (Figure 1E). No significant cleavage products were detected by either antibody. Anti-aSyn antibodies appeared to showed stronger reactivity for aSyn-GFP relative to the untagged form by immunoblot (Figure 1 E and Extended Figure 1). Nonetheless, *Snca* ^*wt/GFP*^ mice expressed both aSyn-GFP and untagged aSyn at ~50% of the amount of aSyn seen in *Snca* ^*GFP/GFP*^ and wt mice, respectively (Figure 1E, lower panel). These results indicate that *Snca-GFP* mice express physiological levels of an aSyn-GFP fusion protein with minimal disruption of endogenous mouse aSyn mRNA and protein levels.

### Regional and subcellular localization of aSyn-GFP in *Snca-GFP* mice is identical to wt animals

Since the endogenous promoter and regulatory elements remain unmodified in *Snca-GFP* mice, we predicted that the regional and subcellular distribution of aSyn-GFP would be comparable between wt and both KI genotypes. Examination of native GFP fluorescence in 40 μm-thick brain and spinal cord sections revealed abundant aSyn-GFP expression particularly in the hippocampus, substantia nigra, striatum, cerebral cortex, globus pallidus, thalamus and olfactory bulb (Figure 2A), mirroring the endogenous pattern previously reported for C57BL/6 mice (Taguchi et al., 2016). Detection by immunofluorescence also showed no obvious differences in regional aSyn distribution between brain sections from *Snca*^*wt/wt*^, *Snca*^*wt/GFP*^ and *Snca*^*GFP/GFP*^ mice (Figure 2B). In addition, GFP-immunoreactivity overlapped completely with that of aSyn in *Snca* ^*wt/GFP*^ mice, further indicating the intact tagged protein is normally expressed nor does it perturbate the distribution of wt aSyn (Figure 2C).

**Figure 2.**
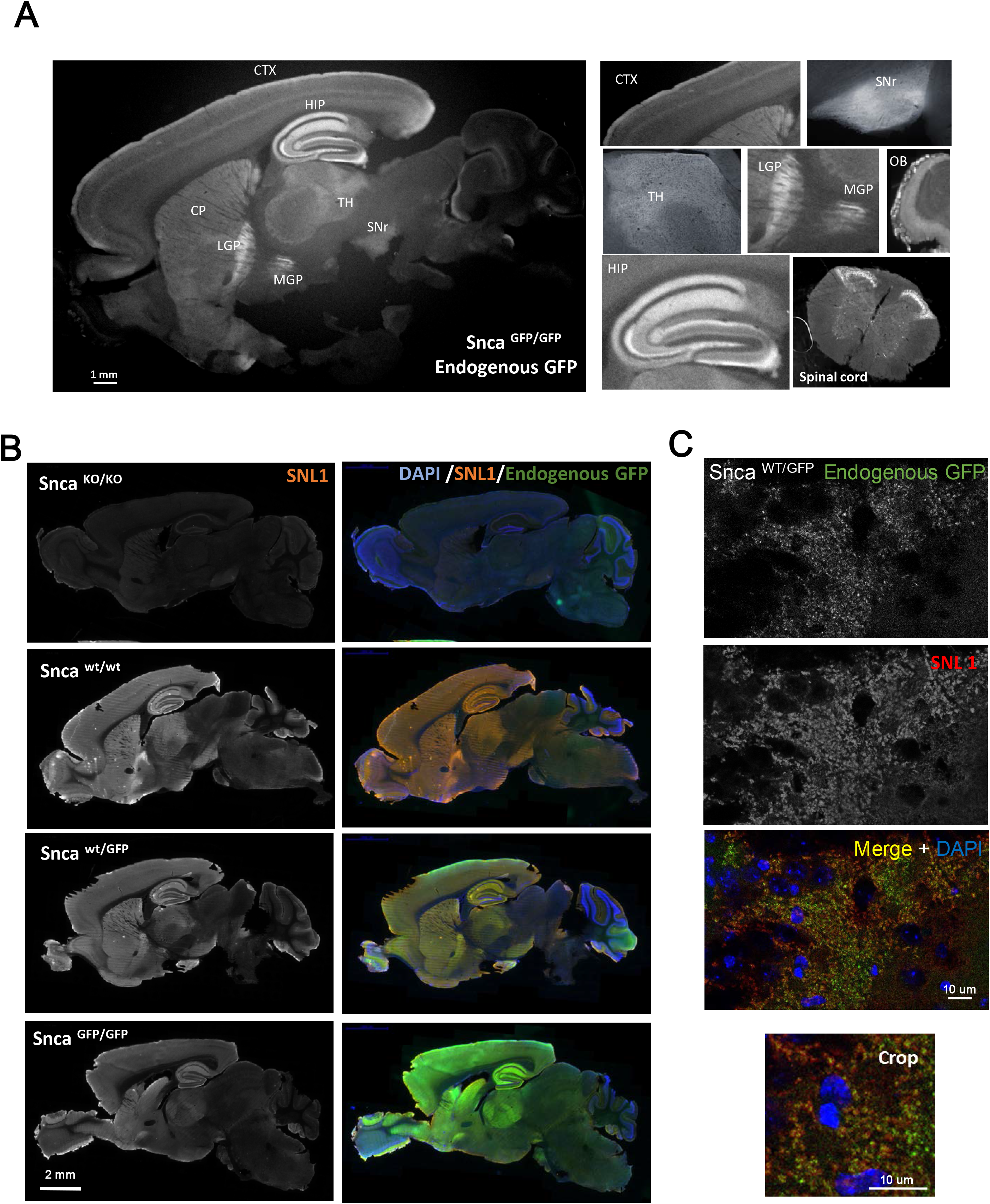
Distribution of aSyn-GFP in brain and spinal cord. **A)** Expression pattern of aSyn-GFP in the brain from a homozygous Snca^GFP/GFP^ animal. Endogenous GFP fluorescence is shown. Individual brain areas are shown on the right. **B)** Immunofluorescence of showing aSyn expression patterns in wt, heterozygous, homozygous Snca^GFP^ and Snca^KO/KO^ mice using SNL1. **C)** Co-localization of aSyn-GFP and endogenous aSyn in the hippocampus of a heterozygous Snca-GFP mouse labeled with a pan-aSyn antibody (SNL1). CTX= Cortex; HIP= Hippocampus; CP= Caudate Putamen; LGP= Lateral Globus Pallidus; MGP= Medial Globus Pallidus; TH= Thalamus; SNr= Substantia Nigra pars reticulata; OB= Olfactory bulb; SI= Substantia Innominata.

Co-immunostaining of GFP with markers for different CNS cell types showed that aSyn-GFP is expressed by neurons but not detectable in astrocytes, oligodendrocytes or microglial cells (Figure 3A). Within neurons, aSyn-GFP co-localized with glutamate vesicular transporter (VGLUT; Figure 3B), a presynaptic marker, consistent with the known enrichment of aSyn in presynaptic vesicles in excitatory neurons (Burre, 2015; Maroteaux et al., 1988). aSyn-GFP was also detectable in brain areas containing non-glutamatergic neurons (e.g., striatum and olfactory bulb), where it co-localized with glutamate decarboxylase (GAD) positive neurons (Figure 3B and data not shown). Together, these results suggest that the *in vivo* distribution of the aSyn-GFP fusion protein closely matches that of wt aSyn at the regional, cellular, and subcellular levels.

**Figure 3.**
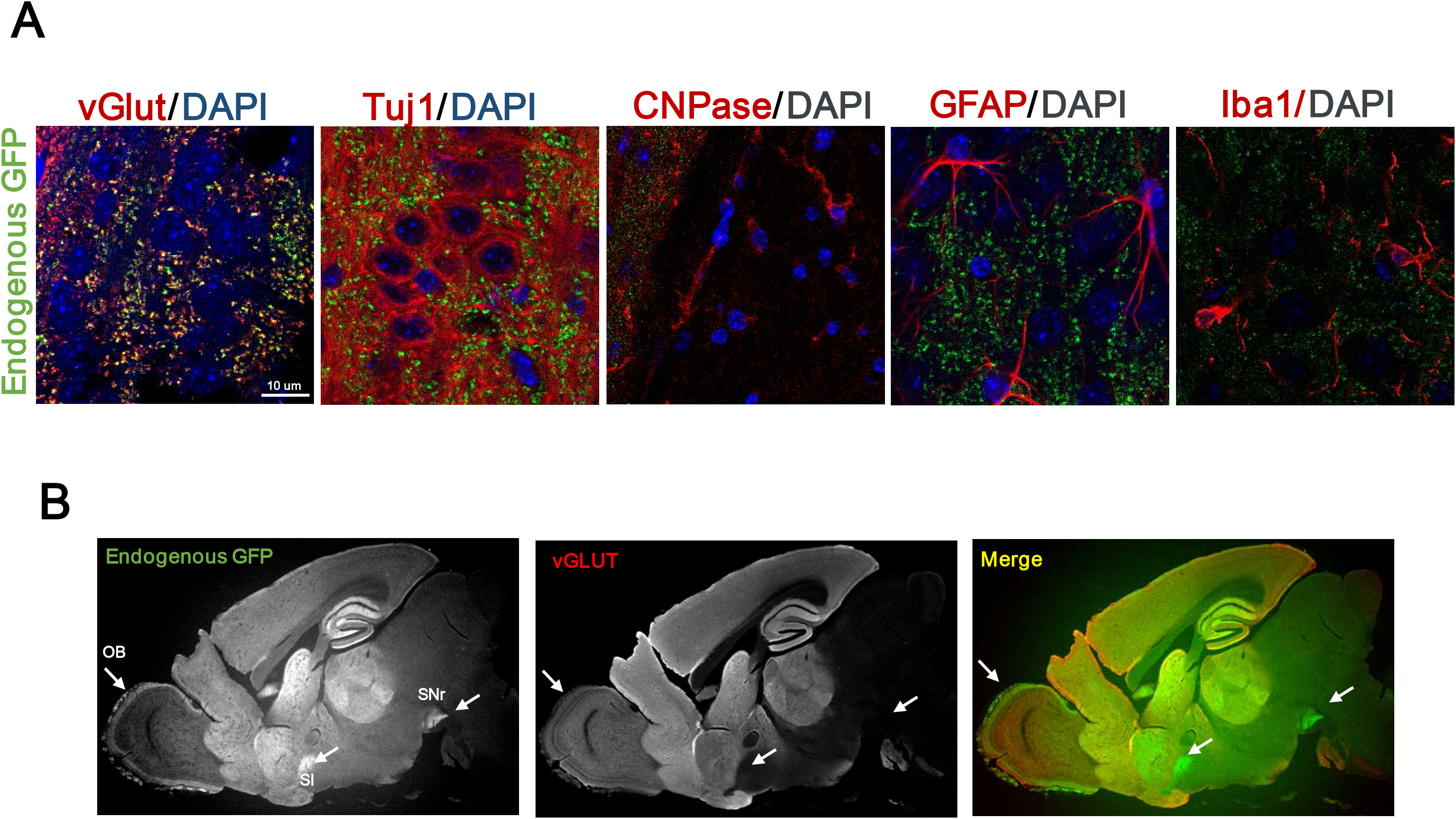
Expression of aSyn-GFP in different CNS cell types. **A)** Co-localization between aSyn-GFP and the indicated cell markers for neurons (VGlut and Tuj1), oligodendrocytes (CNPase), astrocytes (GFAP), and microglia (IBA1) show that the protein is expressed in neurons. **B)** Sagittal brain section of a Snca ^GFP/GFP^ mouse stained with an antibody against vGlut, marker of excitatory presynapses. Arrows indicate brain areas in which aSyn-GFP signal is enriched but vGlut is not present. OB= Olfactory bulb; SI= Substantia Innominata; SNr= Substantia Nigra pars reticulata.

### aSyn-GFP is enriched in presynaptic vesicles in *Snca-GFP* primary neurons

To further determine the utility of these KI mice for studying aSyn pathobiology, we compared the expression levels and subcellular localization of aSyn-GFP in primary neurons derived from *Snca*^*wt/GFP*^, *Snca*^*GPF/GFP*^ and wt mice. Hippocampal and cortical neurons (Figure 4 and data not shown) prepared from postnatal day 0 *Snca*^*wt/GFP*^ and *Snca*^*GPF/GFP*^ mice developed normally in culture, with aSyn-GFP detectable by DIV 7. aSyn-GFP was mainly enriched in vesicles, similar to what is seen in *Snca-GFP* brains (Figure 4A). Both wt aSyn and aSyn-GFP expression in cultured neurons increased with age before plateauing at DIV 10, indicating that the addition of the GFP-tag does not alter the developmentally regulated expression of aSyn in neurons (data not shown). In mature cultures, aSyn-GFP colocalized strongly with Synapsin 1, a marker of synaptic vesicles (Figure 4A). In addition, aSyn and aSyn-GFP strongly overlap in *Snca-GFP*^*wt/GFP*^ neurons, which express equal quantities of both (Figure 4B). Therefore, neurons prepared from Snca-GFP mice exhibit the expected synaptic distribution of aSyn and Synapsin 1.

**Figure 4.**
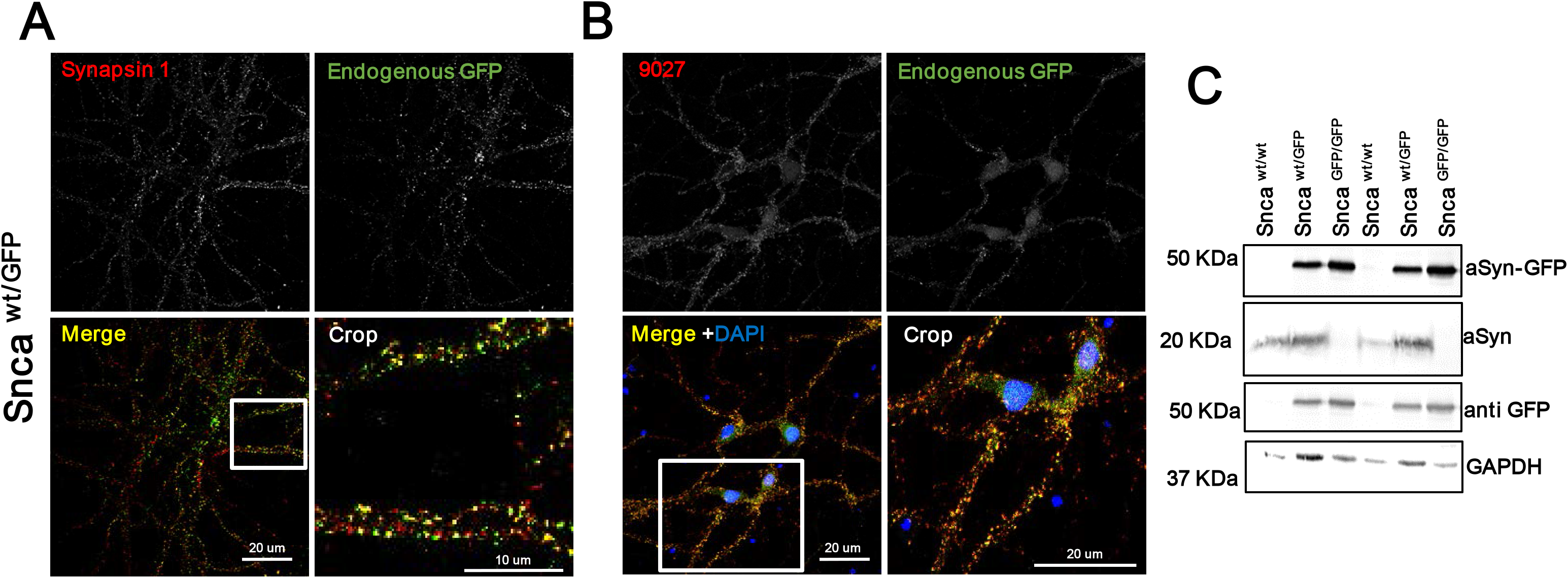
Expression and subcellular localization of aSyn-GFP in primary neurons. Primary hippocampal neurons were derived from newborn pups and kept in culture for 14-21 days. **A)** Co-localization of aSyn-GFP with the presynaptic marker Synapsin1. **B)** Co-localization between endogenous GFP signal and the anti aSyn antibody 9027 in Snca^WT/GFP^ neurons. **C)** Western blot of lysates from neurons of the indicated genotypes using antibodies against aSyn, GFP and GAPDH.

Immunoblot analysis of hippocampal culture lysates with aSyn and GFP antibodies confirmed the expression of a single major species consistent with intact aSyn-GFP at the expected ratios with respect to the wt protein in each genotype (Figure 4C). Taken together, aSyn-GFP maintains normal subcellular expression and distribution without disrupting the morphology of synaptic vesicles.

### aSyn-GFP participates in the synaptic vesicle cycle with kinetics similar to wt aSyn

To further establish that aSyn-GFP has similar functional properties to wt aSyn, we examined its ability to participate in synaptic vesicle cycling in primary hippocampal neurons. In wt neurons, aSyn is dispersed following stimulation with 90 mM KCl and partially repopulates into vesicles after stimulus removal (Fortin et al., 2005). We therefore performed live imaging on DIV18-22 *Snca*^*w/tGFP*^ hippocampal neurons to monitor the redistribution of aSyn-GFP during stimulation and subsequent recovery. Addition of KCl induced a marked decrease in aSyn-GFP intensity in individual vesicles within 2 minutes of treatment (Figure 5A and 5B), in agreement with the dynamics previously reported for wt aSyn (Fortin et al., 2005). Neurons were also fixed and immunostained for Synapsin 1, which is known to disperse following stimulation and redistribute back to synaptic vesicles shortly after the stimulus end (Chi et al., 2001). Unlike Synapsin 1, aSyn-GFP did not fully re-establish its vesicular localization after a 15-minute recovery period following KCl treatment Figure 5C). These results further indicate that aSyn-GFP is functionally similar to its wt counterpart.

**Figure 5.**
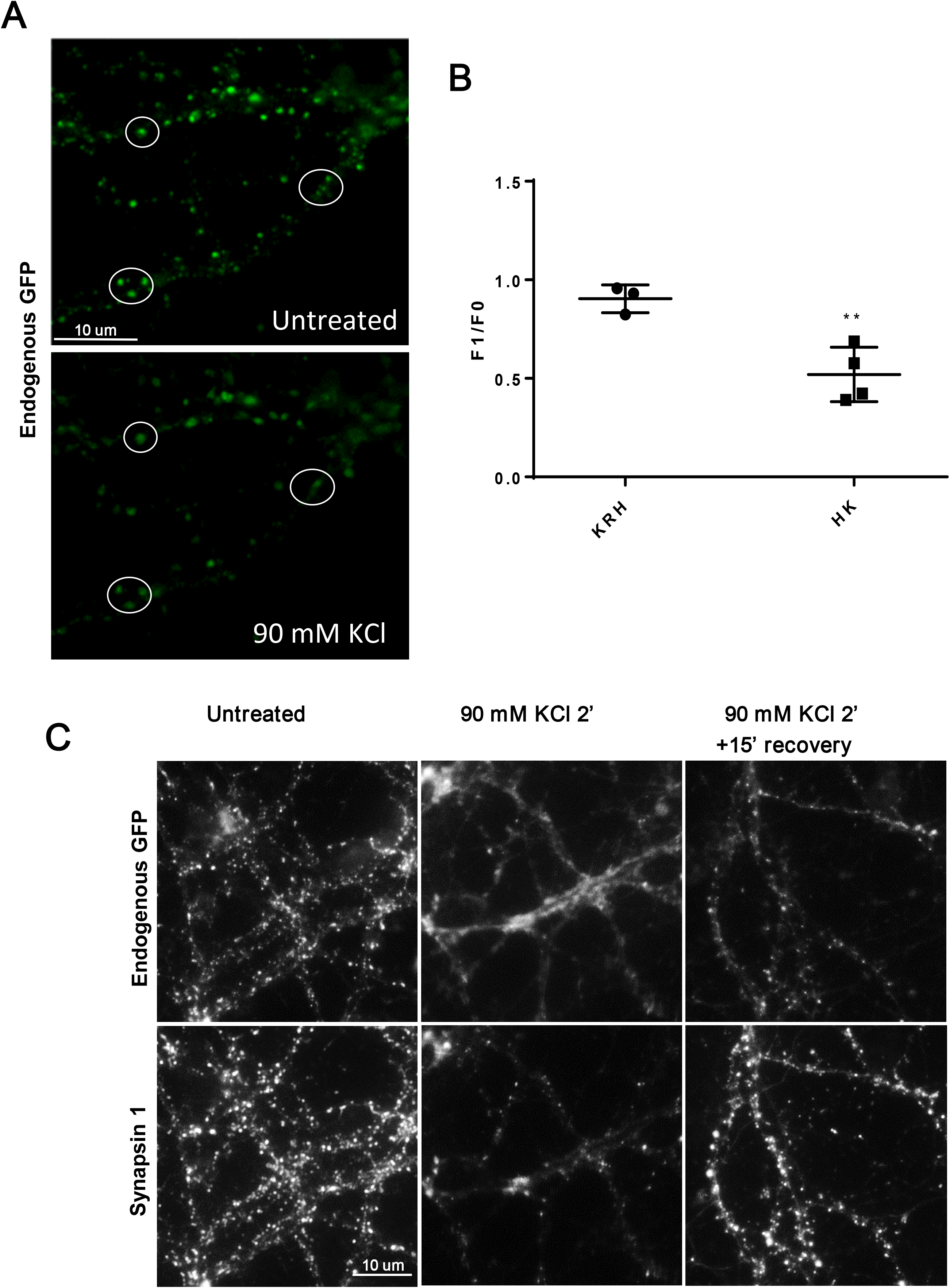
aSyn-GFP behaves similarly to wt aSyn in the synaptic vesicle cycle. **A)** Heterozygous SGKI primary hippocampal neurons were imaged before and after stimulation with 90 mM KCl. **B)** Quantification of fluorescence intensity of GFP positive vesicles before (F0) and following (F1) treatment with control (KRH, Krebs Ringer HEPES) or stimulation solution (HK, 90 mM KCl). The data is expressed as the ratio F1/F0 (mean + SEM, 2 subfields per dish, more than 70 vesicles per dish, N=3-4 dishes from 2-3 independent cultures). Asterisks indicate statistical significance (P= 0.0061) as determined by two tailed unpaired Student’s t-test with Welch’s correction. **C)** Snca^WT/GFP^ neurons were fixed before, after a 2 minute exposure to 90 mM KCl, or after 15 minutes recovery from high potassium stimulation and co-stained with an anti-Synapsin 1 antibody.

### aSyn-GFP is detectable in multiple peripheral cell types

Since expression of aSyn-GFP in *Snca-GFP* mice remains under the control of the endogenous promoter and regulatory elements, we predicted that its distribution in peripheral organs would also parallel that of the wt protein. We therefore surveyed aSyn-GFP expression in a variety of organs across multiple anatomical regions. As previously reported for wt aSyn (Barbour et al., 2008), GFP fluorescence was prominent in blood (primarily erythrocytes; Figure 6A) and bone marrow (data not shown). Endogenous GFP fluorescence in blood was also detectable in *Snca*^*GFP/GFP*^ mice by spectrophotometry (Figure 6B) and confirmed by immunoblotting in lysates prepared from sedimented red blood cells (Figure 6C). We also detected GFP fluorescence in unfixed colon samples (Figure 6D, left panel) in which aSyn-GFP could be found in neurons of the myenteric plexus. Although the native GFP signal was lost after fixation, expression of the fusion protein was confirmed in this tissue using an anti-GFP antibody (Figure 6D, right panels). In contrast, GFP fluorescence in most other tested tissues was indistinguishable from wt mice suggesting levels below our detection limit, consistent with previous studies.

**Figure 6.**
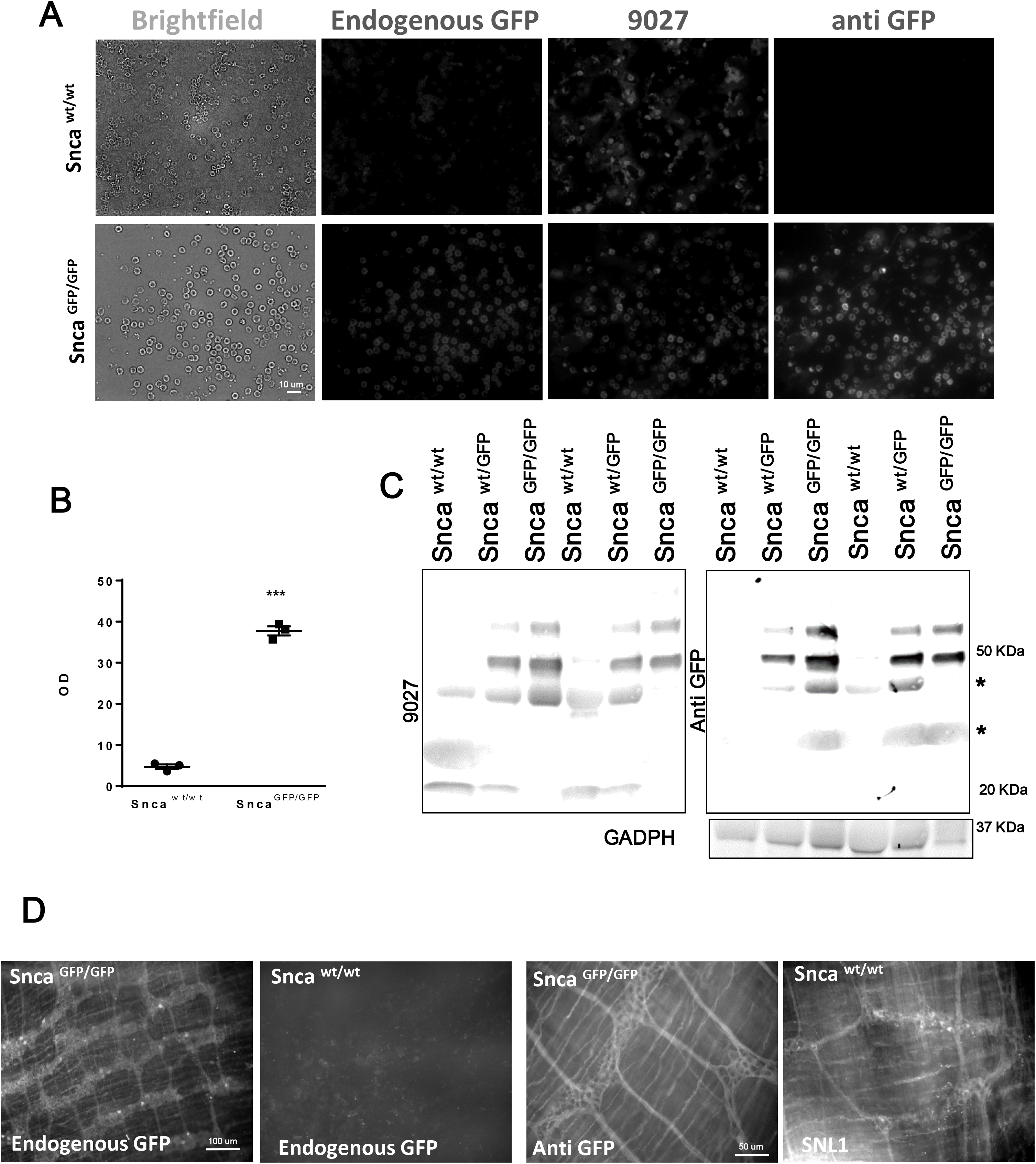
Expression of aSyn-GFP in peripheral cells. **A)** Total blood from Snca ^WT/WT^ or Snca ^GFP/GFP^ mice were fixed on a coverglass and aSyn-GFP was detected using either endogenous GFP fluorescence, an anti-GFP antibody, or anti-aSyn antibody (9027). **B)** Spectrophotometric reading of GFP signal from lysed total blood from either Snca ^WT/WT^ or Snca^GFP/GFP^ mice. Data is shown as mean ±SEM and statistical significance is indicated by asterisks (two-tailed unpaired t-test with Welch’s correction; N=3 animals/genotype, P<0.0001). **C)** Western blot confirmation of the expression of aSyn-GFP in red blood cells. Asterisks denote non-specific bands. **D)** aSyn-GFP in the myenteric plexus of the colon of unfixed whole mounts from Snca^GFP/GFP^ mice detected using endogenous fluorescence (left) or with an anti-GFP antibody in fixed samples (right). Wt aSyn was detected in fixed samples using an anti-aSyn antibody (SNL-1) (N=3 per genotype).

### GFP-tag partially inhibits but does not abolish aSyn fibril assembly

The normal distribution and functioning of aSyn-GFP in *Snca-GFP* mice led us to examine its utility for investigating the pathobiology of aSyn, specifically its formation into amyloid fibrils that accumulate within intracellular Lewy pathology found in PD and DLB (Arima et al., 1998; Takahashi and Wakabayashi, 2001). Since previous studies showed that the presence of a GFP tag can alter the rate of aSyn aggregation (Afitska et al., 2017), we tested whether recombinant aSyn-GFP can self-assemble into fibrils under conditions where wt aSyn readily polymerizes. Independent reactions containing either recombinant wt aSyn or aSyn-GFP monomer were incubated with agitation as previously described (Luk et al., 2016). Within 6h of the start of the reaction, the majority of wt aSyn had converted to insoluble species which accounted for nearly all aSyn by 8h (Figures 7A-C). In parallel reactions, aSyn-GFP also accumulated in the insoluble fraction over time, but with significantly delayed kinetics (Figure 7A-C). When equimolar concentrations of untagged aSyn and aSyn-GFP were combined, both forms aggregated at a similar rate that fell between that of each individual protein (Figure 7C). Inspection of the products from these reactions by electron microscopy (Figure 7D), revealed the presence of filamentous structures compatible with those previously reported for aSyn (Luk et al., 2009). Of note, aSyn-GFP fibrils assembled in vitro are able to induce aSyn pathology in wt and Snca-GFP neurons ((Karpowicz et al., 2017) and data not shown). Taken together, aSyn-GFP is fibril assembly-competent, albeit with slower kinetics, which may be mitigated in the presence of untagged aSyn monomer.

**Figure 7.**
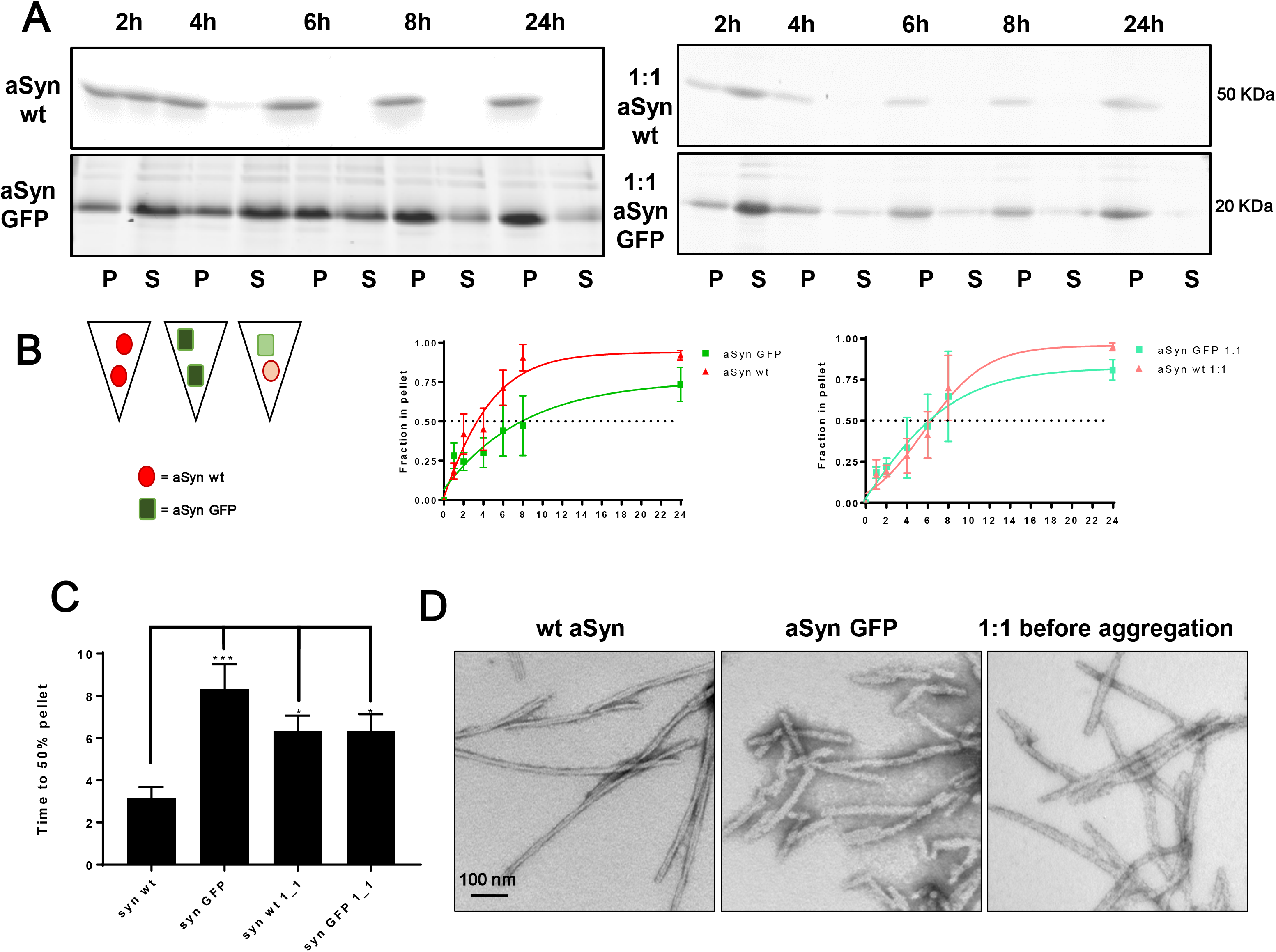
aSyn-GFP is assembly-competent in vitro and in primary neurons. **A)** Recombinant wt, aSyn-GFP or a mixture of both (1:1) were agitated for the indicated times at 37°C. Insoluble pellet (P) and soluble supernatant (S) fractions were separated after ultracentrifugation and visualized by Coomassie Blue staining after SDS-PAGE. **B)** Quantification of data in A, expressed as the relative percentage of protein in the pellet fraction over time. Data (N = 3-5 per time point, from 2 different batches of monomers) were fitted to a sigmoidal curve. **C)** Time to 50% pellet calculated after curve fitting (Mean ± SEM) to a sigmoidal curve. C) Time to 50% pellet (Absolute EC50) was calculated after fitting the data (Mean+ SEM) to a sigmoidal curve. One-Way Anova with Tukey’s post-test was used to analyze statistical significance. * indicates P<0.05; *** P=0.002. **D)** Transmission electron microscopy images of negatively stained fibrils from the indicated monomer(s) after 7 days of shaking show fibril formation from all the monomers tested (N=4).

### Incorporation of aSyn-GFP into pathological aggregates following PFF-seeding in *S*nca-GFP neurons

We and others have previously demonstrated that aSyn fibrils internalized by neurons can template the conversion of endogenously expressed aSyn into fibrillar forms that accumulate as Lewy-like inclusions (Volpicelli-Daley et al., 2011). We therefore determined whether primary hippocampal neurons from *Snca-GFP* mice are permissive to such pathological seeding following exposure to aSyn PFFs and incorporated aSyn-GFP into insoluble intraneuronal inclusions. Neurons from *Snca*^*wt/GFP*^ mice developed inclusions resembling Lewy neurites and Lewy bodies when exposed to mouse wt PFFs, although the level of pathology, as measured by pSer129 aSyn (pSyn) immunostaining, was reduced relative to similarly treated wt neurons (Figure 8 A and B). In agreement with our *in vitro* data showing that the rate of aggregation is reduced when only aSyn-GFP is present, pathology formation at the same time point was further reduced in *Snca*^*GFP/GFP*^ neurons. Nevertheless, pSyn was still detectable in a small proportion (~0.005% of wt) of PFF-exposed *Snca*^*GFP/GFP*^ neurons (Figures 8A and 8B).

**Figure 8.**
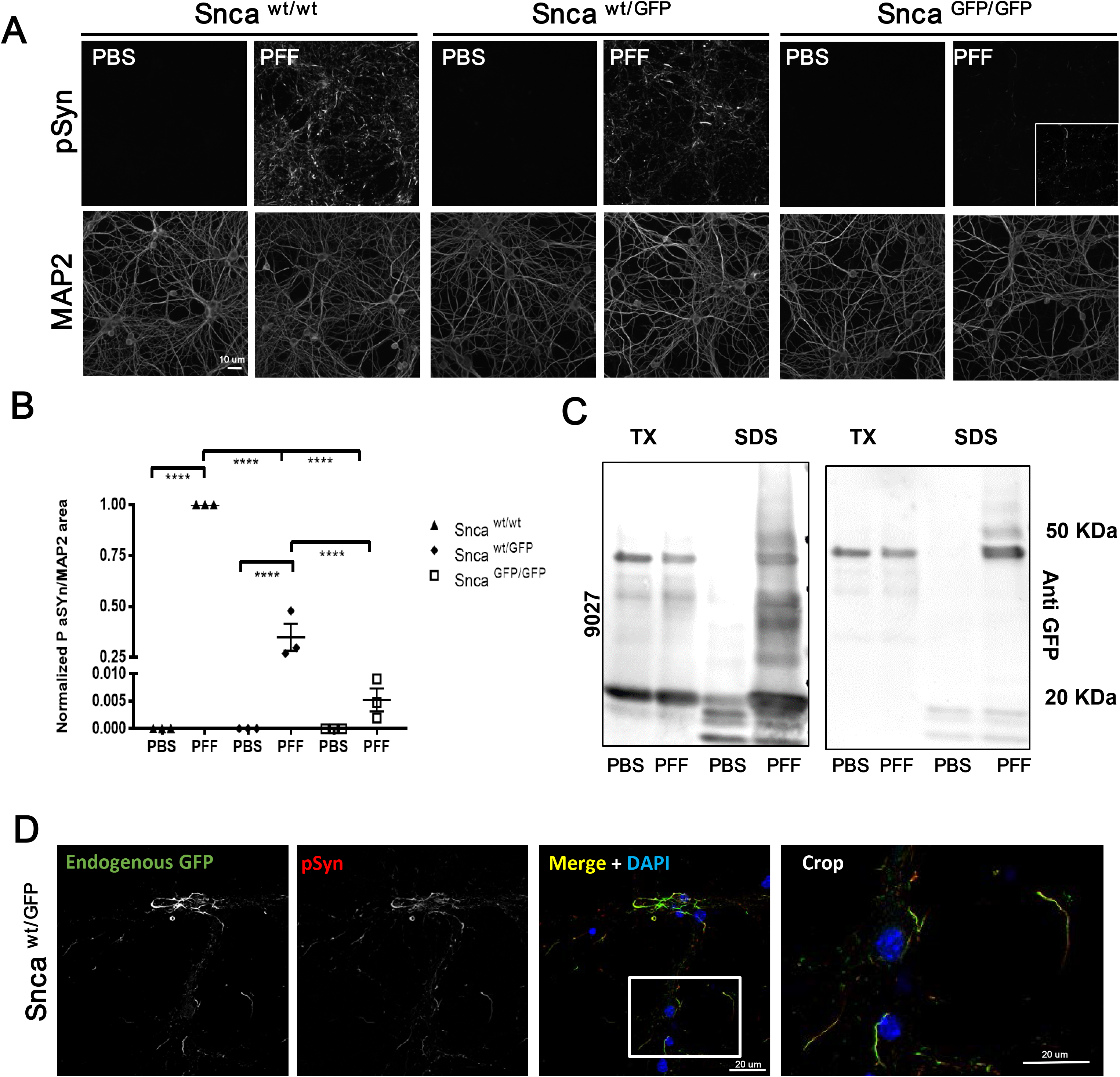
aSyn GFP forms LB like aggregates in primary neurons. **A)** Primary hippocampal neurons were treated with mouse wt aSyn PFFs at 10 DIV and phosphorylated synuclein (pSyn) was detected by immunofluorescence after 2 additional weeks. MAP2 was used to confirm presence of neurons and to normalize pSyn levels. In each experiment, cultures from all 3 genotypes were present on individual plates and subjected to same treatments and handling. 3-4 wells/batch of culture were used and normalized to wt cultures (N = 3 from 3 independent cultures). **B)** Quantification of data in A normalized to wt cultures (9-12 wells from N=3 independent cultures/genotype; Mean ± SEM). Asterisks indicate statistical significance following Two-Way ANOVA with Sidak’s multiple comparison test. P<0,0001. **C)** Sequential extraction in 1% Triton X-100 and 2% SDS of PBS or PFFs treated Snca ^WT/GFP^ neurons indicates that aSyn-GFP shifts to the insoluble fraction upon PFFs exposure. **D)** Immunostaining of PFF treated Snca^WT/GFP^ neurons with an anti-pSyn antibody (81A) shows nearly complete overlap with endogenous GFP fluorescence.

In both *Snca*^*wt/GFP*^ and *Snca*^*GFP/GFP*^ neurons, co-labeling with antibodies to pSyn and GFP revealed near-complete co-localization within inclusions, indicating that aSyn-GFP was uniformly incorporated (Figure 8D and data not shown). Biochemical analysis of Triton-X100 insoluble proteins from these cultures also confirmed that a majority of aSyn-GFP shifted to this fraction after PFF treatment (Figure 8C).

We further determined whether aSyn-GFP can also undergo pathological conversion *in vivo* by using intracranial injection to target PFFs into the mouse brain, a model that allows the induction and propagation of aSyn pathology in presence of wt levels of this protein. For this, we selected the hippocampus due to previous reports (Luna et al., 2018; Nouraei et al., 2018) and our present data showing high levels of aSyn/aSyn-GFP expression in this region. In agreement with this, injection of wt mouse aSyn PFFs into the hippocampus induced the formation of pSyn-positive inclusions within 30 days post injection (Figures 9A and B). Pathology levels were highest in wt mice, followed by *Snca*^*wt/GFP*^ animals, with very low pathology observed in *Snca*^*GFP/GFP*^ mice (Figures 9A and C). Similar results were obtained by injecting PFFs into the dorsal striatum (Extended Figure 9). In both injection paradigms, pSyn highly co-localized to endogenous GFP fluorescence (Figure 9 B and Extended Figure 9), indicating that the fusion protein is incorporated in aSyn aggregates similarly to what we observed in cultured neurons (Figure 8 A and B). Of note and consistent with the injection data, Snca-GFP mice do not develop pathology during normal aging (data not shown) marking a notable difference with previous models overexpressing GFP-tagged aSyn (Hansen et al., 2013).

**Figure 9.**
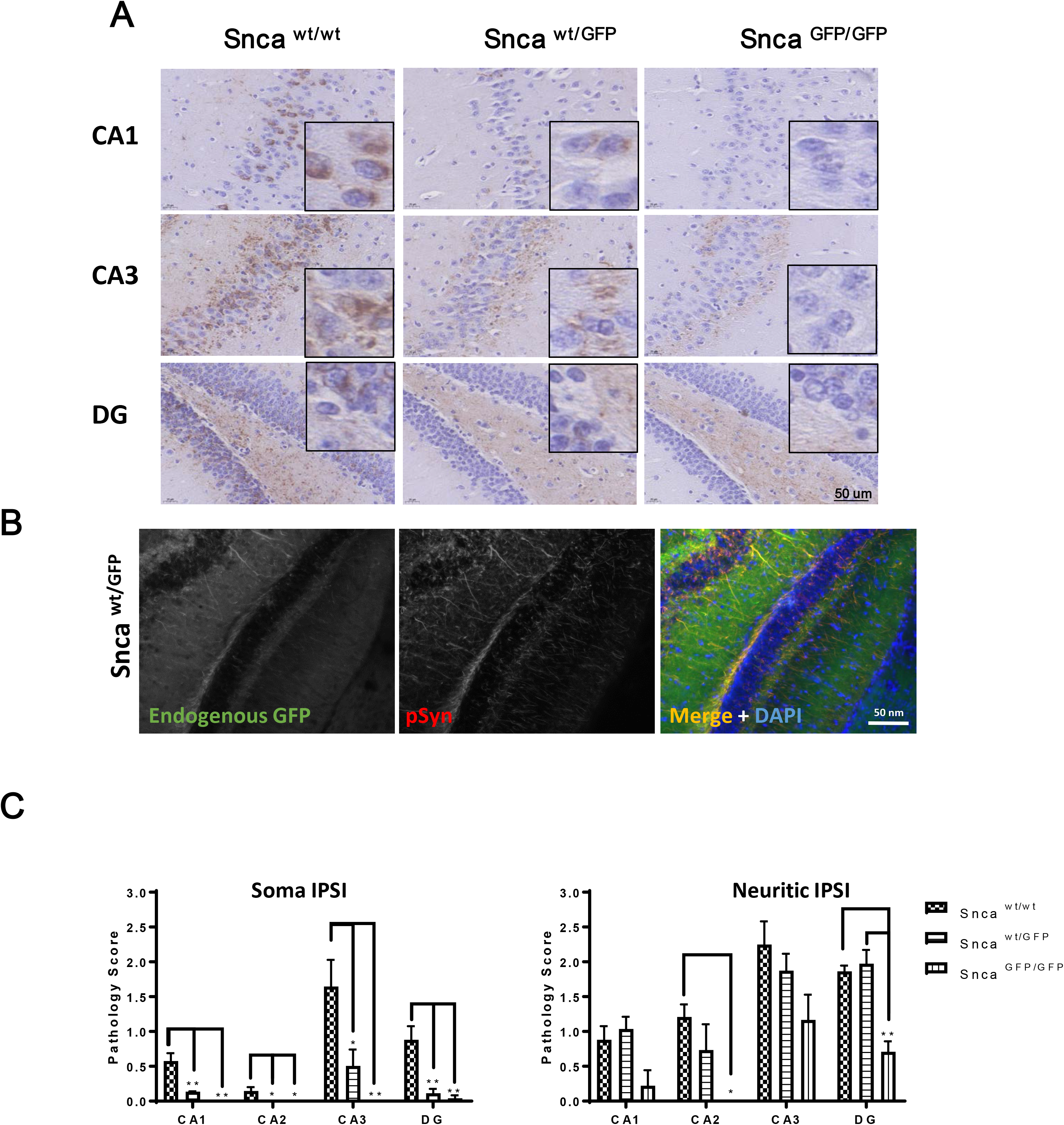
Intracranial injection of aSyn PFFs induces formation of intraneuronal pathology in Snca ^WT/GFP^ mice. 5 ug of recombinant mouse aSyn PFFs were injected into the hippocampus of Snca^WT/WT^, Snca^WT/GFP^, or Snca^GFP/GFP^ mice. Brains were analyzed 30 days post injection (dpi) by immunohistochemistry **(A)** and immunofluorescence **(B)** using an anti pSyn (81A) antibody. **C)** Quantification of the data in A. Plots indicate mean ± SEM. Asterisks indicate statistical significance following One-Way Anova (brain area) with Tukey’s post-hoc test. (* P<0.05, ** P<0.005). CA= Cornu Ammonus; DG= Dentate Gyrus.

## Discussion

Clinical and experimental evidence implicate aSyn in multiple neurodegenerative disorders. However, neither the normal function and regulation of aSyn, nor its role in disease initiation and progression, are fully understood. Given the dynamic nature of these biological processes, tools that enable the direct visualization of aSyn would facilitate efforts to address these fundamental questions. Moreover, the majority of individuals with synucleinopathy carry neither aSyn mutations nor overexpress aSyn to any measureable extent. To this end, we generated a aSyn-GFP knock-in (KI) mouse line in which endogenous aSyn protein is fused to GFP via its carboxy-terminal with the goal of combining the utility of a genetic-encoded fluorescent tag while preserving the protein’s natural distribution throughout the body.

Our data here demonstrate that both heterozygous (*Snca*^*wt/GFP*^) and homozygous (*Snca*^*GFP/GFP*^) mice express aSyn-GFP in a pattern that is indistinguishable from aSyn protein in wt animals across multiple tissues including, but not limited to, the central and enteric nervous systems, erythrocytes, and bone marrow. Within the brain, where aSyn is highly enriched, total aSyn mRNA and protein levels were comparable between wt and KI animals while expression levels of aSyn-GFP were directly proportional to gene dosage. In contrast to some previous reports, aSyn-GFP appeared to be expressed primarily in its intact form and minimal levels of degraded products were detected (McLean et al., 2001). We speculate that expression of aSyn-GFP at endogenous levels and the inclusion of the short linker between aSyn and GFP may contribute to this stability. When examined *in vivo* or in cultured neurons derived from KI mice, aSyn-GFP is detectable in all neurons and is correctly localized to a subset of synaptic vesicles, while the complete co-localization between native and GFP-tagged aSyn in *Snca*^*wt/GFP*^ mice suggests that aSyn-GFP does not perturb the expression or localization of its wt counterpart. Indeed, the addition of the GFP-tag appears to have a minimal effect on a complex aspect of aSyn function (i.e., participation in synaptic vesicle cycling) as both wt and tagged aSyn disperse similarly after synaptic stimulation, in agreement with previous studies (Fortin et al., 2005; Unni et al., 2010).

The above characteristics distinguish this KI line from previously reported mammalian models that employ a genetically encoded GFP tag to label aSyn. In these, the tagged sequence corresponds to wt human aSyn with expression regulated by either a non-*Snca* promoter (e.g. PDGFβ) (Rockenstein et al., 2005) or downstream of the mouse *Snca* promoter by means of a bacterial artificial chromosome (Hansen et al., 2013). Although robust neuronal expression was achieved in both examples, the distribution and levels of tagged aSyn did not precisely match that of aSyn in the non-transgenic host and aSyn was also ectopically expressed in additional cell types and regions. Interestingly, GFP-tagged aSyn in these mouse lines also accumulate as lysosome-associated inclusions or undergo phosphorylation at Ser129, a marker of Lewy body and Lewy neurite pathology in human synucleinopathies, with aging. A possible explanation for such differences is that the previously described lines were selected based on high expression levels of the transgene, among other criteria. Total aSyn levels in these mice may have been further amplified by maintaining these mice on a wildtype genetic background without deletion of the endogenous *Snca* locus. In contrast, the localization and levels of aSyn-GFP in *Snca*^*wt/GFP*^ and *Snca*^*GFP/GFP*^ mice matched that of wt mice and we did not observe any redistribution or modification of aSyn-GFP in mice up to 2 years of age. We believe these features make aSyn-GFP KI mice particularly amendable for investigating aSyn trafficking and action, a crucial but understudied area of aSyn biology, especially given that these processes are highly sensitive to aSyn levels (Eguchi et al., 2017; Scott and Roy, 2012).

An added advantage is that aSyn-GFP is normally distributed throughout peripheral organs such as neurons in the gastro-intestinal tract and red blood cells in this line. As we demonstrate, this expression also permits rapid quantification of aSyn levels without the need for additional sample processing (e.g. immunostaining). It is anticipated that this will further enable the validation of endogenous aSyn expression in other peripheral tissues, especially those where aSyn is in low abundance, and where detection using immunohistochemistry alone has provided equivocal results. Primary cells derived from aSyn-GFP KI mice are also a resource for investigating the aSyn biology at the cellular level.

In addition to physiological functioning, our work here demonstrates that *Snca*^*wt/GFP*^ and *Snca*^*GFP/GFP*^ neurons can also serve as permissive cellular host for pathological seeding by misfolded aSyn species. Specifically, recombinant aSyn PFFs induced GFP-positive Lewy-like pathology in cultured hippocampal neurons when introduced into the culture media or in multiple CNS regions when PFFs are stereotaxically injected into either dorsal striatum or hippocampus. Importantly, fibril-induced pathology in *Snca*^*wt/GFP*^ and *Snca*^*GFP/GFP*^ was distributed in the same brain regions as in wt mice, albeit with different densities, suggesting a similar spreading process within neuroanatomical pathways. The intact aSyn-GFP moiety represented the major species in these intraneuronal inclusions, confirming that this pathology is formed predominantly by aSyn-GFP derived from the neuronal pool and is consistent with previous studies showing that neurons overexpressing aSyn-GFP support fibril-induced pathology formation (Osterberg et al., 2015; Volpicelli-Daley et al., 2014). Interestingly, the proportion of untagged aSyn and aSyn-GFP found in the intracellular inclusions in PFF-treated *Snca*^*wt/GFP*^ neurons were similar, providing further support that the tagged protein is converted under a similar process.

Although our data clearly shows that aSyn-GFP can polymerize into fibrils and can be recruited into Lewy-like pathology in neurons, the kinetics of this process is altered by the presence of the GFP tag compared to wt untagged aSyn both *in vitro* and *in vivo*. These observations are in agreement with previous reports and underline the importance of the C-terminus domain and protein flexibility in facilitating conversion to a pathological conformation (Afitska et al., 2017; Bertoncini et al., 2005; McLean et al., 2001) although the constraints brought on by the sterically larger GFP moiety does not abrogate seeding activity. Indeed, aSyn-GFP monomers readily assemble under the same *in vitro* conditions as untagged aSyn and can be incorporated into fibrils at near stoichiometric levels in presence of an equimolar concentration of wt aSyn monomer, with aggregation kinetics that are intermediate relative to wt aSyn or aSyn-GFP alone. In line with these *in vitro* observations, fibril-induced pathological seeding in neurons and *in vivo* also show a similar pattern with pathology formation in neurons from wt mice forming inclusions the most rapidly, followed by *Snca*^*wt/GFP*^ and then *Snca*^*GFP/GFP*^ mice.

In summary, we have generated a novel *in vivo* resource for studying multiple aspects of aSyn function under physiological and disease-like conditions without genetic overexpression. Snca-GFP mice provide the opportunity to concomitantly track and measure soluble and pathological forms of the protein across relevant tissues. We anticipate that future studies leveraging these animals (e.g., by crossbreeding with other genetic models of disease) should provide additional insights into aSyn biology.

## Methods

### Animals

All housing, breeding, and procedures were performed according to the NIH Guide for the Care and Use of Experimental Animals and approved by the University of Pennsylvania Institutional Animal Care and Use Committee. Animals were anesthetized with a mix of Ketamine: 100 mg/kg; Xylazine: 10 mg/kg; Acepromazine: 0.5 mg/kg before performing transcardiac perfusion with PBS + heparin (2 USP/ml). The *Snca*-GFP KI line was created by homologous recombination: a synthetic mouse Snca exon 6 with a short linker (the primary sequence around the linker being …EPEA-KL-MVSKG…) was inserted directly after the last amino acid of the mouse *Snca* coding sequence, followed by the enhanced GFP coding sequence and the remainder of the SNCA 3’UTR (Figure 1A). A synthetic DNA construct (Blueheron Inc.) consisting of ~200 nt of the genomic Snca sequences upstream of exon 6 and all of exon 6 (with the above linker and eGFP sequence inserted directly after the C-terminus of Snca) was cloned directly downstream of the floxed Neo cassette of PL452 (NCI, Frederick). A ~500 nt long Snca upstream arm of homology was added upstream of the Neo cassette. This construct was introduced by BAC recombineering into pL253 (NCI, Frederick) harboring ~12 kB of the genomic SNCA region containing at its center SNCA exon 6. The resulting targeting vector contained a 14183 nt long Snca allele with integrated floxed Neo cassette as well as Snca exon 6 containing KL – eGFP downstream of the Snca protein. The targeting vector was linearized with NotI and introduced into V6.5 ES cells by electroporation. After geneticin selection ES clones were screened by Southern blotting (HindIII digestion) and 5 positive clones were found and further validated by a second round of Southern blotting (PvuII digestion) using a different probe (see Figure 1 A). The positive clones generated a wt ~12 kB and a mutant ~9kB PvuII fragment. Positive clones were subjected to chromosome counting. Clone 2B9, which had the correct number of chromosomes, was injected into C57/B6 mouse blastocysts. Heterozygous and homozygous mice were obtained and confirmed by PCR using primers encompassing the fusion region (Exon 6) that amplify a longer fragment when the GFP sequence is present (See Figure 1A and B) and resulted in phenotypically normal mice until 24 months of age. No effect on survival was observed (data not shown). Snca-GFP mice were bred with a Cre-recombinase expressing line (Jackson line 019099) to remove the neomycin cassette. Animals negative for the CRE gene were identified by PCR and selected for subsequent breeding (See Figure 1 B) although no differences were observed between Neo+ or - mice. C57BL/6 J were purchased from Jackson (JAX 000664). *Snca*^−/−^ mice (Abeliovich et al., 2000) were maintained on a C3H background.

### Antibodies

See table 2

**Table 1.**
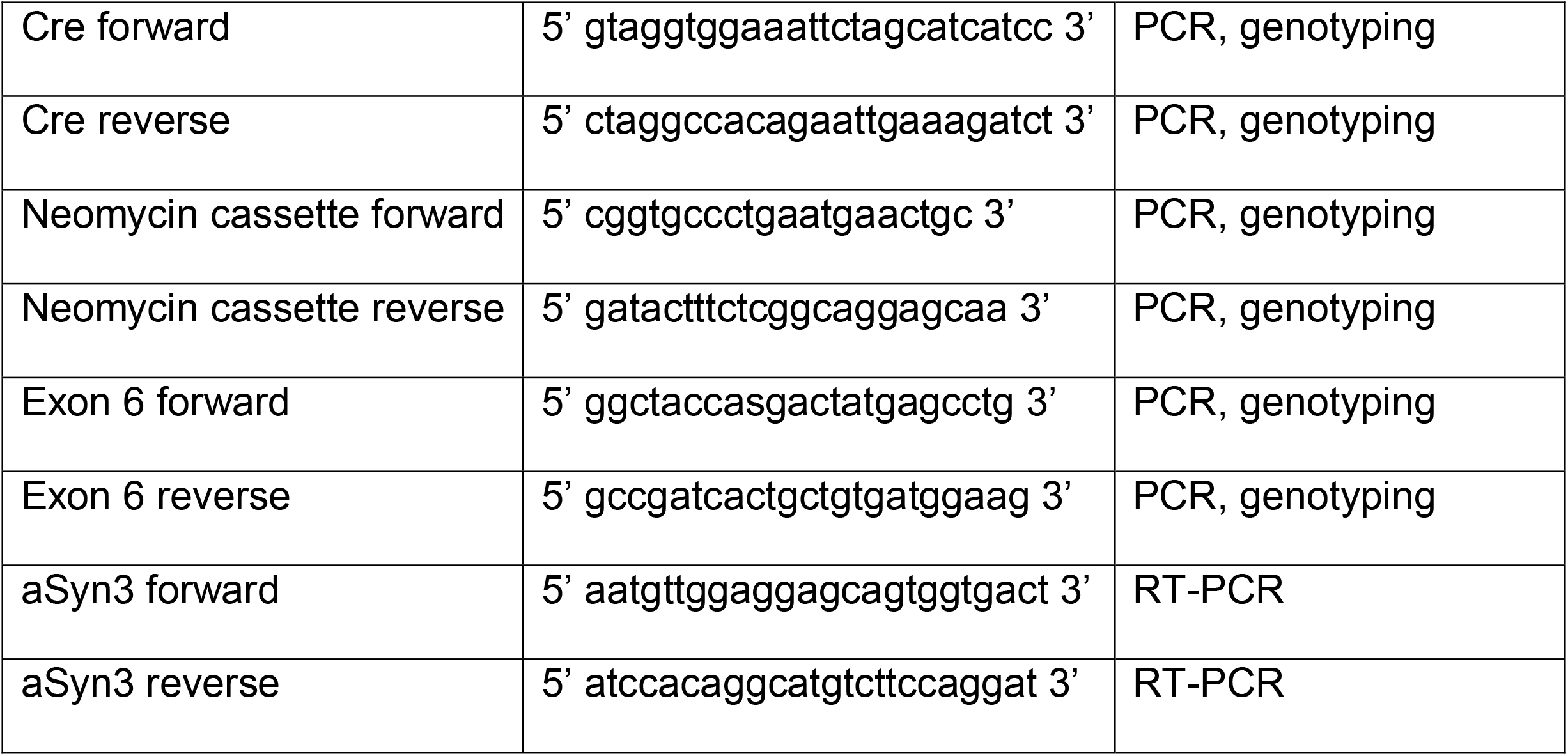
PCR primers used in this study

**Table 2.** Antibodies used in this study

| Antibody | Antigen | Host | Use and Dilution | Source |
| --- | --- | --- | --- | --- |
| 9027 | alpha-synuclein | Mouse | WB1:10K<br>IF 1:5K | CNDR (Covell et al., 2017) |
| SNL-1 | alpha-synuclein | Rabbit | WB, IF<br>1:1K | CNDR (Giasson et al., 2002) |
| GFP | Green Fluorescence Protein | Rabbit | WB 1:1K | Novus Biologicals |
| GFP | Green Fluorescence Protein | Chicken | IF 1:2K | Aves labs |
| GFAP | Glial fibrillary acidic protein | Rabbit | IF 1:1K | DAKO |
| MAP-2<br>(17028) | Microtubule Associated Protein 2 | Rabbit | IF 1:2K | CNDR (Volpicelli-Daley et al., 2011) |
| Iba-1 | Macrophage/Microglia-Specific Calcium-Binding Protein (Iba1) | Rabbit | IF 1:2K | DAKO |
| GAPDH<br>(6C5) | Glyceraldehyde-3-phosphate dehydrogenase | Mouse | WB<br>1:1.5 K | Advanced Immunochemical |
| Synapsin-1<br>AB1543 | Synapsin 1 a and b | Rabbit | IF 1:1 K | Millipore |
| Tuj-1 | Neuronal Class III Beta-Tubulin (TUJ1) | Mouse | IF 1:1K | BioLegend |
| vGlut<br>135304 | VGLUT 1 (vesicular glutamate transporter 1 | Guinea Pig | IF 1:2 K | Synaptic Systems |
| 81A | a-synuclein phospho S129 | Mouse | IF, IHC<br>1:8K | CNDR (Waxman and Giasson, 2008) |
| MJFR13 | a-synuclein phospho S129 | Rabbit | WB 1:1 K | Abcam |

### Reagents and Chemicals

All reagents were purchased from Fisher and all chemicals were purchased from Sigma unless otherwise indicated

### Recombinant aSyn monomers and PFFs

Recombinant mouse aSyn and aSyn-GFP were produced as previously described (Karpowicz et al., 2017; Luk et al., 2012). In brief, plasmids encoding the 2 proteins were introduced in *E. coli* using heat shock and proteins were purified following lysis using size exclusion and anion exchange chromatography. Purified protein (monomer) was concentrated to 5 mg/ml for aSyn wt and 15 mg/ml for aSyn GFP (360 uM) and kept frozen until use. Recombinant PFFs (mouse wt aSyn) were produced by shaking 500 ul of recombinant aSyn monomer (5 mg/ml, 360 uM) at 37 C for 7 days. Fibril formation was confirmed by sedimentation assay (see specific section). Right before their use, PFFs are diluted in PBS and sonicated for 10 cycles (1 sec ON, 30 sec OFF, high intensity) in a bath sonicator at 10°C (BioRuptor; Diagenode).

### Electron Microscopy

Wildtype aSyn (2.5 mg/ml, 180 uM), aSyn-GFP (7.5 ug/ml, 180 uM), or wt aSyn + aSynGFP (assembled as 1:1 monomer mixture) were diluted in PBS, absorbed onto Formvar/carbon film-coated copper grids (Electron Microscopy Sciences), washed twice with water and negatively stained with 0.4-2% uranyl acetate. Grids were visualized with a Jeol 1010 transmission electron microscope (Peabody).

### Sedimentation assay

Monomers were thawed and ultracentrifuged at 100,000 g for 30 minutes at RT. They were then diluted to 2.5 (wt aSyn, 180 uM) and 7.5 mg/ml (aSynGFP, 180 uM) in PBS. 100 ul of asyn, aSyn GFP or aSyn + aSyn GFP (1:1) were aliquoted in Eppendorf tubes and shaken at 1,000 rpm at 37°C for 24h. 6 ul of each reaction was collected at the indicated time points, diluted in 60 ul of PBS and layered on a 25% sucrose cushion before undergoing ultracentrifugation (30 mins at 100,000 × *g*). Supernatant and pellet fractions were separated. Pellets were resuspended in 120 ul of 12.5% sucrose. 25 ul of 5X SDS sample buffer were added to each tube and 30 ul was loaded on 15% acrylamide gels for SDS PAGE. Separated proteins were then stained with Coomassie Blue ON and destained with 10% isopropanol, 10 % acetic acid and water before image acquisition with an infrared scanner (LiCor Odyssey).

### Sequential extraction

Neurons were washed twice with PBS and lysed in TBS (50 mM TRIS, 150 mM NaCl, pH=7.6) + 1% Triton X-100. Lysates were sonicated for 1 second ON, 30 seconds OFF, medium intensity in a bath sonicator at 10 C (BioRuptor; Diagenode) and then rotated at 4 C for 20 minutes. Lysates were then spun at 100,000 × *g* for 30 mins at 4°C, supernatants were removed and labelled as TX-100 soluble fraction. The pellet was washed once with 1% TX-100 TBS, sonicated and ultracentrifuged. Supernatants were discarded and pellets dissolved in 1/10 volume of TBS, 2% SDS, sonicated and ultracentrifuged. Supernatants were collected as SDS soluble fraction and pellets were discarded. Protein content was measured by BCA in the TX-100 soluble fraction. Equal amount of total proteins from TX-100 soluble fractions were separated by SDS-PAGE (4-20% gradient gel). An equal volume of the SDS soluble fraction was loaded alongside. Proteins were detected by western blot with the indicated antibodies.

### Immunofluorescence

*Primary neurons:* Neurons were fixed at the indicated DIV with warm 4% paraformaldehyde+ 4% sucrose for 15 minutes at RT. Cells were permeabilized and non-specific binding sites blocked using PBS containing 0.1% TX-100, 3% BSA, 3% FBS for 20 minutes. Primary antibodies were added to cells for 1 h, followed by 3 washes with PBS and 1 h incubation with the appropriate secondary antibody (1:1 K). Cells were washed 3 times with PBS, once in water and mounted using Fluoromount-G.

*40 um brain sections and intestinal whole mounts:* sections were permeabilized and blocked in PBS containing 10% FBS, 3% BSA, 0.5% TX-100 for 1 hour at RT, incubated with primary antibodies ON at RT, washed 3 times with PBS, incubated with secondary antibodies for 2 hours at RT, washed and mounted using Fluoromount-G (brain sections) or 1:1 PBS/glycerol (intestine).

*6 um sections:* sections were produced and treated as described in the immuonohystochemistry section with the modification that after the ON incubation with primary antibodies, sections were incubated with a fluorescently labelled secondary antibody for 2 hours at RT and mounted using Flouromount-G.

Images were captured on a Nikon Ds-Qi1Mc digital camera attached to a Nikon Eclipse Ni microscope (6 um sections), a Leica Confocal SP8 for colocalization studies (primary neurons and 40 um sections) or using InCell Analyzer 2200 (GE Healthcare) when using 96 well plates). Analysis was performed using Fiji or Developer software (GE Healthcare, 96 well plates).

### Immunocytochemistry

PBS-perfused mouse brains were post fixed in ethanol (70% in 150 mM NaCl) ON at 4°C, cut in 3 mm slabs, and embedded in paraffin. Tissue was then sectioned at 6 micrometers using a microtome and applied on glass slides. Before staining, sections were de-paraffinized and rehydrated (xylene, 95%, 90% and 75% ethanol). Sections were blocked in TBS containing 3% FBS and 2% BSA for 1 hour at RT, incubated with primary antibodies at 4 C ON, with biotinylated secondary antibodies for 1 hour at RT (Vector Laboratories), horse radish peroxidase conjugated streptavidin for 1 hour at RT (Vector Laboratories), and signal was revealed using DAB peroxidase substrate products (dark brown, Vector Laboratiories). Sections were counterstained with hematoxylin for 1 min and mounted using Cytoseal Mounting Media. Images were captured on a Nikon Ds-Qi1Mc digital camera attached to a Nikon Eclipse Ni microscope or using Lamina Scanner (PerkinElmer; 20X objective).

### Synaptic vesicle cycling

Hippocampal neurons were plated on IBIDI dishes and kept in culture for 18-21 days. Cells were then incubated in Krebs Ringer HEPES buffer (5 mM KCl, 140 mM NaCl, 10 mM HEPES, 10 mM Glucose, 2.6 mM CaCl_2_, 1.3 mM MgCl_2_) for 15 minutes before starting the imaging sessions. Images were acquired with a Leica microscope and a 40X air objective. Cells were imaged (every 3-5 seconds) in KRH solution for 2-3 minutes to establish a baseline signal. The media was then switched to high potassium (HK; 90 mM KCl, 55 mM NaCl, 10 mM HEPES, 10 mM Glucose, 2.6 mM CaCl_2_, 1.3 mM MgCl_2_) or KRH solution for 2 minutes. Individual vesicles were fragmented using ImageJ/FIJI and intensities before and after stimulation were determined as ratios after subtracting background signal and adjusting for image drift (Lazarenko et al., 2018).

For fixed samples, cells on coverslips were washed and incubated in KHR for 15 minutes at RT and then either fixed immediately before treatment, incubated with HK for 2 mins before fixation (HK), or washed for an additional 15 mins using HRK before fixation (HK+15’ recovery). Cells were then processed for immunostaining as described above.

### Western Blot

Cells were lysed in lyses buffer (0.5% Triton X-100, 0.5% deoxycholic acid, 10 mM TRIS, 100 mM NaCl, pH=8.0) with phosphatase and protease inhibitors. Nuclei and debris were removed by centrifugation (5 mins at 1,000 × *g*). Total protein content was determined by BCA and equal amounts of total protein were separated on 4-20% gradient gels. Proteins were then transferred to 0.22 um nitrocellulose membranes (1 hour at 100 V at 4 C). Membranes were blocked in 7.5 % BSA and incubated with the indicated primary antibodies ON followed by appropriate secondary antibodies (LiCor) for 1 hour at RT. Image acquisition was performed using a Licor Scanner and image analysis performed using ImageJ/FIJI (NIH).

### Primary neurons and PFF transduction

Primary hippocampal neurons were derived from P0-P2 SGKI pups. Hippocampi were dissected in MEM 10 mM HEPES 1% P/S, digested with 1 mg/ml Papain for 30 minutes and dissociated with a 1 ml tip in MEM 10% FBS, 2mM GlutaMax, Sucrose, 1% Penicillin/Streptomycin. Cells were centrifuged at 1K RPM for 4 minutes, resuspended in MEM/2%B27, 2mM GlutaMax and plated at the density of 4.5×10^4/cm2.

PFF transduction was conducted at 10 DIV by adding 2 ug/ml (96 well plate and coverslips) or 10 ug/ml (biochemistry) of PFFs diluted in PBS and sonicated with a BioRuptor (Diagenode) bath sonicator for 10 cycles of 30 seconds on, 30 seconds off, high setting). Cells were processed 14 days post transduction (DPT).

### Intracerebral injection of PFFs

2.5 ul of 2 mg/ml of sonicated mouse wt pre formed fibrils (PFFs) were stereotaxically injected in the hippocampus (coordinates: − 2.5 mm relative to Bregma; 2 mm from midline; 2.4 mm beneath the skull; Figure 9) of male mice of the indicated genotype (2-4 months of age, 3 mice/genotype; (Zhang et al., 2019). Mice were perfused 30 days post-surgery. Additionally, as shown in Extended Figure 9, PFFs were stereotaxically injected into the ventral striatum (AP: +0.2 mm Bregma, lateral: 2.0 mm from midline, depth: 3.6 mm beneath the dura), dorsal striatum (AP: +0.2 mm, lateral: 2.0 mm, depth: 2.6 mm), and overlaying cortex (AP: +0.2 mm, lateral: 2.0 mm, depth: 0.8 mm) of mice (females) of the indicated genotype and mice were perfused at 30 (1 mouse/genotype, data not shown) or 90 days post injection (1 mouse/genotype).

### RT-PCR

Mice were perfused, brain removed in conditions that minimize RNA degradation and half hemispheres were frozen for further processing. Frozen tissue was thawed and immediately homogenized in 2 ml of RTL buffer + 2mM DTT (Qiagen). mRNA was extracted from 300 ul of homogenate using the RNeasy Kit (Qiagen) according to manufacturer’s instructions. 200-800 ug of mRNA were converted to single strand cDNA with SuperScript™ III First-Strand Synthesis System (Thermo Fisher) using random hexamers and the protocol detailed by the vendor. Brain derived cDNA and primers targeting mouse a-Syn (see table) were used in Syber Green (Thermo Fisher) Real Time PCR reactions monitored by a 7500 Fast Real Time PCR system (Applied Biosystems). Actin and SNAP 25 were used as internal controls (see table). Data is expressed as fold change over the wild type genotype.

### Measurements of GFP signal in mouse blood

Total blood was collected in EDTA containing tubes by cardiac puncture in deep terminal anesthesia and right before transcardial perfusion (see specific section). 25 ul of blood were lysed by addition of 225 ul of lysis buffer and 5 ul of lysate were diluted in 190 ul of water and analyzed for fluorescence using a spectrophotometer (excitation: 488 nm; emission: 530 nm).

### Experimental designs and Statistical analysis

Details for each experiment and statistical analysis are described in the Figure Legends. Statistical analysis was performed using GraphPrism (v7) or GraphPrism (v8 for figure 7C).

## Acknowledgments

We thank Victoria Kehm for technical assistance, Myrna Dominick, EunRan Sun and Chi Li for their help with RT-PCR experiments, and Hakeem O. Lawal for critical reading of the manuscript. We also thank all members at the Center for Neurodegenerative Disease Research (CNDR) for continuous discussions. This work was supported by NIH grant NS088322 (KCL).

---

**Extended Figure 1.**
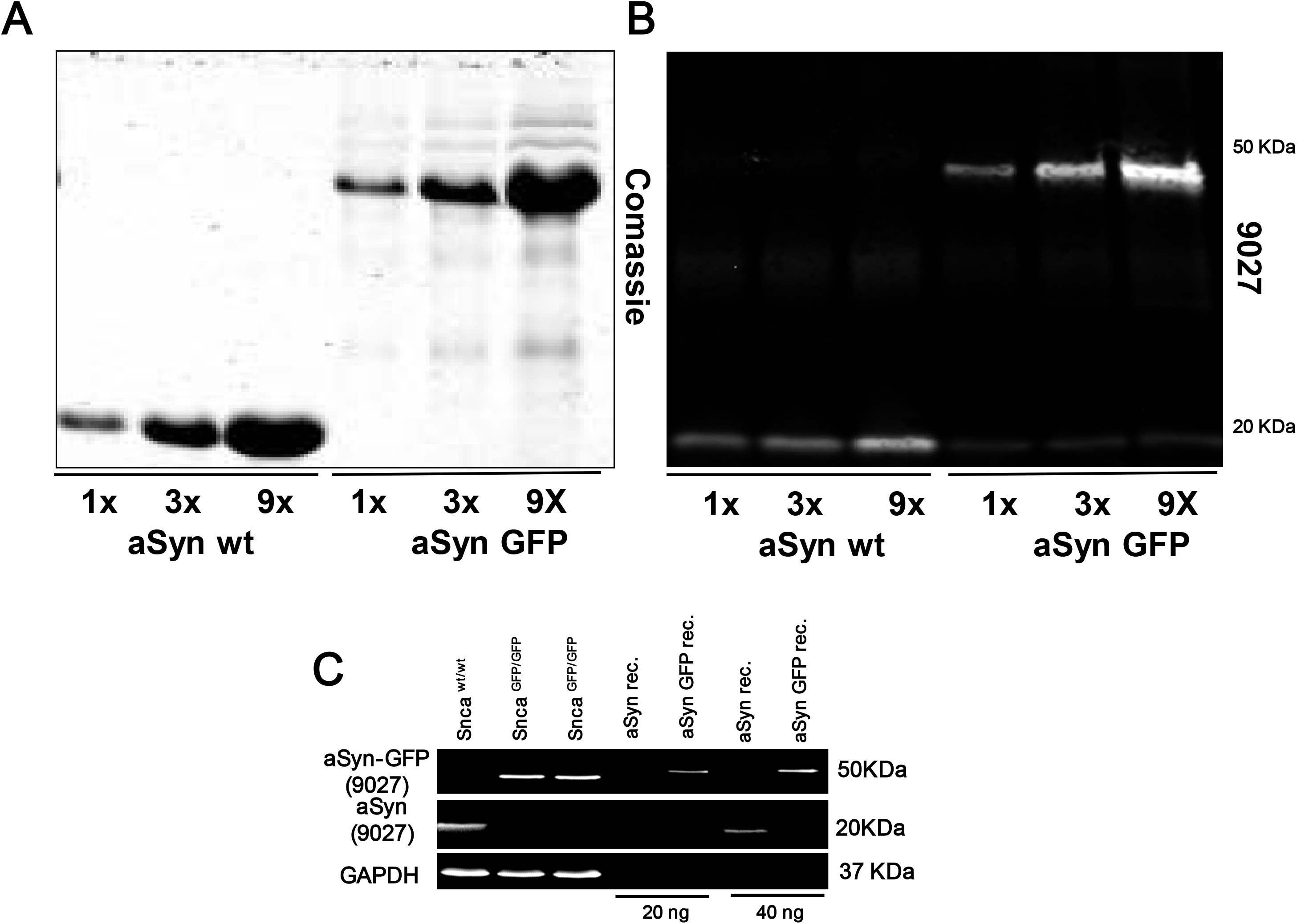
**A)** Coomassie staining and **B)** western blotting with aSyn antibody (Syn9027) of different amounts of recombinant mouse aSyn and aSyn-GFP. **C)** Western blot of brain homogenates of the indicated genotype loaded along with the indicated amount of recombinant aSyn and aSyn-GFP proteins.

**Extended Figure 9.**
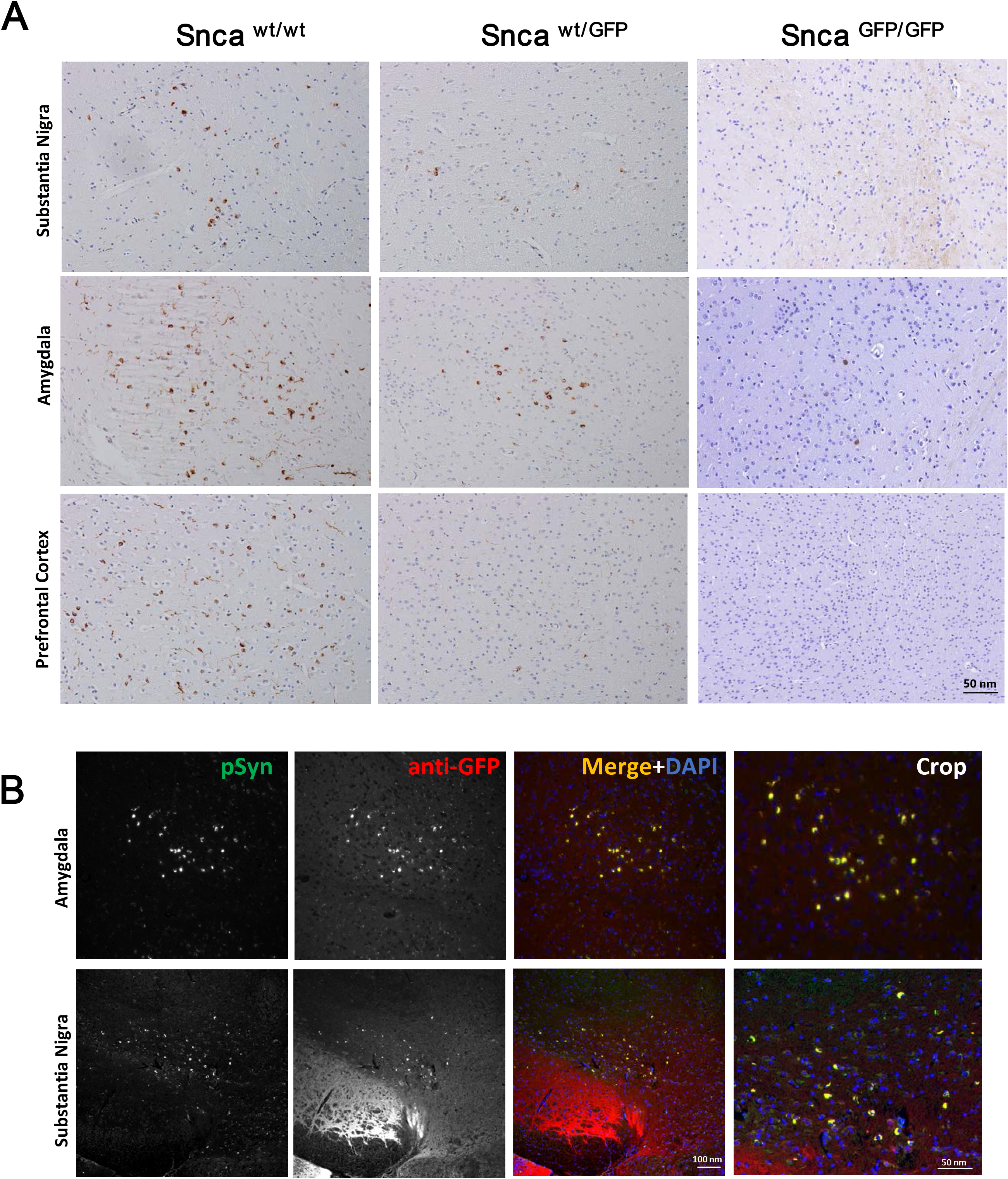
5 ug of recombinant preformed fibrils were injected in the dorsal striatum, ventral striatum and cortex of wt, heterozygous and homozygous Snca^GFP^ mice. Pathology was analyzed at 90 days post injection (dpi) by immunohistochemistry **(A)** and immunofluorescence **(B)** using an antibody (81A) against phosphorylated aSyn (pSyn).

